# Harnessing Escherichia coli’s Dark Genome to Produce Anti-Alzheimer Peptides

**DOI:** 10.1101/2023.06.23.546343

**Authors:** Neeraj Verma, Siddharth Manvati, Pawan K. Dhar

## Abstract

Alzheimer’s disease (AD) is characterized by progressive neurodegeneration. The critical molecular trigger is believed to be the accumulation of Aβ neurotoxic oligomers. Given the proteolytic processing of Amyloid Precursor Protein (APP) by β-secretase (beta-site APP cleaving enzyme 1, BACE1) as the key step in the building up of Aβ oligomers, BACE inhibitors come with therapeutic prospects of preventing or delaying the onset of Alzheimer’s. To find inhibitory peptides against BACE1, a library of ‘dark peptides’ was constructed from 4400 intergenic DNA sequences of *Escherichia coli*. The sequence level analysis was followed by protein structure predictions, molecular docking, and simulation. Based on bioinformatics analysis, 5 potential peptides were screened for experimental validation. Out of these two peptides were identified as lead molecules based on BACE1 inhibitory activity, followed by FRET inhibitory assay, western blot, and RT-PCR. An 86.7 % drop in BACE1 level was observed in the presence of the ECOI2 peptide. Though encouraging results were obtained from in-silico and in-vitro studies, more work is required to study the efficacy of these peptides in suitable animal models.

## Introduction

Alzheimer’s disease (AD) is a progressive neurodegenerative disorder that affects millions of people worldwide. It is the most common cause of dementia. AD is characterized by the accumulation of amyloid plaques and neurofibrillary tangles in the brain, leading to synaptic loss and neuronal death. The clinical manifestations of AD include memory impairment, language difficulties, visuospatial deficits, behavioral changes, mood disturbances. The diagnosis of AD is based on clinical criteria, neuropsychological testing, cerebrospinal fluid (CSF) analysis and brain imaging.

The β-secretase enzyme, also known as BACE (β-site amyloid precursor protein cleaving enzyme), plays a crucial role in the development of Alzheimer’s disease. The enzyme is responsible for cleaving the amyloid precursor protein (APP) generating β-amyloid peptides that collectively form amyloid plaques found in the brains of Alzheimer’s patients.

Elevated levels of BACE and increased activity of the enzyme have been observed in people suffering with Alzheimer’s disease. Amyloid plaques trigger a series of events resulting in neuronal dysfunction and degeneration. The plaques disrupt communication between neurons, leading to memory loss, cognitive decline, and other symptoms associated with Alzheimer’s disease.

To study the onset of plaques and find therapeutic solutions, recombinant BACE1 has been designed and found to cleave APP producing an β-amyloid in the in-vitro system.

Given the critical role of BACE in the production of β-amyloid peptides, the enzyme has become an important target for developing therapeutic intervention. Several drug candidates have been designed to inhibit BACE and studies have been taken up to clinical trials to reduce the production of β-amyloid and halt the progression of the disease (Huang et al, 2020). Despite rapid advances, an effective solution is still lacking.

In search of novel and effective solutions against Alzheimer’s, we extracted molecules from the dark genome of E. coli. The term ‘dark genome‘ refers to the non-expressing, non-translating, and extinct DNA sequences that can be artificially encoded into functional molecules

This idea originates from our first paper where growth inhibitory proteins were synthesized from intergenic sequences (Dhar et al 2009). Following this initial proof of the concept, several antimicrobial, anticancer, anti-parasitic peptides were made from the non-expressed DNA sequences of *E*.*coli, D*.*melanogaster* and *S*.*cerevisiae*.

The current study was designed to decipher the function of non-expressing DNA-derived peptides and proteins with a hope to find novel molecules against Alzheimer’s.

## Materials & Methods

### Creation of Escherichia coli intergenic sequence-derived Peptide database

We retrieved the 4400 intergenic sequences of E. coli strain K-12 sub strain MG1655, from the EcoCyc database. Using Exapsy sequences were computationally translated. Those sequences that showed stop codon were removed. A total of 835 intergenic sequences in the range between 10-35 amino acids were considered for further studies. Peptides were further characterized based on molecular properties like the length of sequences, molecular weight, Isoelectric point, hydrophobicity, hydrophilicity, net charge and so on.

### In-silico screening and 3D model predictions of peptides for their anti-Alzheimer’s potential

To screen the therapeutic potential of intergenic sequence-derived peptides against Alzheimer’s disease peptides were screened against Neuropedia and WALTZ-DB, an anti-Alzheimer’s peptide database using the smith waterman algorithm. The cell-penetrating properties were virtually identified using the CellPPD tool (Gautam *et al*, 2013 and 2015). Models were evaluated using a five-fold cross-validation technique. Peptides were modelled into the 3D structure using Pepfold 3 Mobyle server http://bioserv.rpbs.univ-parisdiderot.fr/services/PEP-FOLD3/

### Cloning of BACE1 and APP

From the human cell line, total RNA was isolated and processed for c-DNA synthesis. The BACE1 and APP genes were amplified from the c-DNA and cloned using pcDNA3.1 and pLVX-Puro vectors respectively using the gateway cloning method to generate BACE1-pcDNA and APP-pLVX constructs for expression. The constructs were sequenced to confirm their construction.

### Development of Human Neuro-blastoma Cell Lines (SH-SY5Y) for expression of BACE1 and APP

The human neuroblastoma cell line SH-SY5Y was purchased from the National Centre for Cell Science (NCCS). Cells were grown in DMEM/Ham’s F-12 (1:1), supplemented with 1X Penstrep and 10% Fetal Bovine Serum (FBS). Cells were maintained at 37°C in a humidified atmosphere containing 5% CO2. SH-SY5Y and stably transfected with plasmids carrying BACE1 and APP genes cloned into mammalian expression vector pcDNA3.1 and pLVX-puro and expressed under the control of the CMV and PGK promoters. The cells were plated at the 6.25 × 10^5^ cells/cm^2^ density in 6-well culture plates (Corning) and each well was transfected with 4 μg plasmid DNA using Lipofectamine 3000 (Invitrogen) as a transfection reagent. The transfected cells were selected with 40 μg/ml G418 for BACE1 expression and 1mg/ml of puromycin in the case of APP. The expression profile of BACE1 and APP gene was confirmed by western blotting.

### Confirmation of Expression by Western Blotting

SH-SY5Y cells were lysed in buffer containing 50 mM Tris-HCl pH 7.4, 150 mM NaCl, 1% Triton X-100, 1 mM EDTA, 0.1% SDS, 1 mM phenylmethylsulfonyl fluoride (PMSF) and 1 X protease inhibitor cocktail (PIC). The cells were incubated with a lysis buffer for 10 minutes on ice before scraping the cells from the dishes. Cell lysates were centrifuged at 13000 rpm for 20 minutes at 4°C. The protein content in the supernatants was measured using the Bradford Protein Assay (Biorad). Samples, containing 50 μg protein, were resolved in 8% SDS-PAGE using Tris-acetate SDS running buffer. The proteins were transferred to nitrocellulose membranes using iBlot™ gel transfer stacks (Invitrogen). Membranes were blocked in PBS with 0.05% Tween 20 containing 3% non-fat dry milk for 1 hour at room temperature.

Mouse anti BACE1 antibody (Santa Cruz Biotechnology) was diluted at 1:1000 and the GAPDH antibody (Santa Cruz Biotechnology) at 1:10000 in 1% skim milk and incubation was carried out at 4°C overnight.

Horseradish-peroxidase (HRP) conjugated anti-mouse secondary antibody (Santa Cruz Biotechnology) was incubated for 1 hour at RT in 1% skimmed milk at the dilution of 1:1000. Blots were developed using the Super Signal^®^ detection system (Pierce). Image J software was used to quantify the blot band density, and the sample loading was normalized to GAPDH.

### Peptide synthesis

The peptides were chemically synthesized (Biocell corporation) at a >99 % purification

### Cell viability assay

Cells (3 × 10^4^ cells/ml) were plated in 96-well plates and allowed to grow for 24 hours. Cells were treated with different peptide concentrations for 24 hours. Cell viability was assessed using the MTT assay. Here, 5 mg/mL MTT (Amresco, USA) was added, and the cells were incubated for 4 hours at 37°C. Afterwards, 150 μL DMSO was added to each well, and the optical density (OD) was measured using a microplate reader (Thermo Labsystems Multiskan MK3, USA) with a 630-nm filter. Cell viability was calculated as relative cell viability = (OD treated − OD blank) × 100%/ (OD control − OD blank).

### BACE1 activity assay based on fluorescence resonance energy transfer (FRET)

To experimentally validate predictions and evaluate the potency of synthetic BACE1 inhibitory peptides, BACE1 activity was measured using a cell-free fluorescence resonance energy transfer (FRET) assay. The test is based on a fluorescence resonance energy transfer (FRET) technique in which the fluorescence signal is enhanced after BACE1 cleaves the substrate. This assay was performed according to BACE1 Activity Detection Kit protocols (Sigma, St. Louis, MO). Briefly, the BACE1 enzyme was prepared in the Fluorescent Assay Buffer to a concentration of approximately 0.3 units/μL and substrates are diluted in the same buffer to a concentration of 50 μM. The peptides were also diluted into different concentrations in DMSO. This assay was performed in 96-well microplates using 100 μl, which consisted of 20 μl of substrate working solution, 2 μl of BACE1 working solution, and 5 μl of different peptide concentrations. The reaction was allowed to proceed for four hours in the dark under a lid at 37°C and was terminated by a stop solution. A Varioskan LUX multimode microplate reader (Thermo Scientific) using 320 nm excitation and 405 nm emission wavelengths, measured the product fluorescence before and after the reaction. The percentage of inhibition was calculated using the following equation: 100-(IFi/IFo×100), where IFi and IFo represent the BACE1 fluorescence intensities in the presence and absence of inhibitor, respectively. Inhibition curves were created by graphing the percentage of inhibition versus the inhibitor concentration. Beta secretase inhibitor IV (C_31_H_38_N_4_O_5_S) has been used as a positive control in this experiment.

### In-vivo expression study of BACE1 and APP proteins by western blotting

A change in APP and BACE1 protein expression levels in the presence of ECOI2-treated SH-SY5Y-APP-BACE1 cells was studied using western blot. Confluent SH-SY5Y-APP-BACE1 cells were treated with different concentrations of 100 nM, 250 nM, 500 nM, 750 nM and 1000 nM of ECOI 2 for 24 hrs. Cells were harvested, and total protein was isolated with the RIPA lysis method. The mixture of proteins containing 120 Kda (APP) and 56 Kda (BACE1) was resolved by SDS PAGE followed by the transfer of proteins to the nitrocellulose membrane.

The APP and BACE1 protein levels were determined by APP-specific monoclonal antibodies and BACE1 protein-specific antibodies. The band intensity was measured by densitometric analysis using Image J software.

The effect of the ECOI2 over the 24 hours was observed in terms of fold change expression. All treatment groups containing 100 nM, 250 nM, 500 nM, 750 nM and 1000 nM concentrations of the ECOI2 yield a significant change in protein level (Figure 9). The experiments were executed in triplicate to verify APP and BACE1 protein levels vis-à-vis the ECOI2 concentrations. GAPDH, a housekeeping protein, was used as a reference to validate protein concentration changes.

### Gene expression study by RT-PCR

To determine how ECOI 2 reduced the accumulation of Aβ1-40 and Aβ1-42 in SH-SY5Y-APP-BACE1 cells, real-time PCR (qPCR) was performed to observe changes in relative mRNA concentrations of amyloid-beta hypothesis-related genes.

The SH-SY5Y-APP-BACE1 cells were treated with 1 μg ECOI 2 for 12 hours cultured for 24 hours. The Aβ metabolism genes were studied, including APP, BACE1, Presenilin 1 (PS1), Presenilin 2 (PS2), Insulin-degrading enzyme (IDE), apolipoprotein E (APOE), low-density lipoprotein related receptor (LRP1), and low-density lipoprotein receptor (LDLR).

## Results

From a total of 835 intergenic sequence-derived peptides against Alzheimer’s disease, 5 peptides (ECOI1-5) were selected based on high smith waterman scores and cell-penetrating properties (CPP). The secondary structure was predicted by the Mobyle server (Lamiable A et al 2016). Docking study of 5 lead peptides against BACE 1. ECOI 1-5 peptides were docked against the BACE1(PDB ID: 1FKN) by DADDOCK easy server (G.C.P van Zundert et al. 2015). Based on the DADDOCK score (figure 3) and the high affinity of peptide toward the target molecule was considered for wet-lab experimental validation

**Fig 1:**
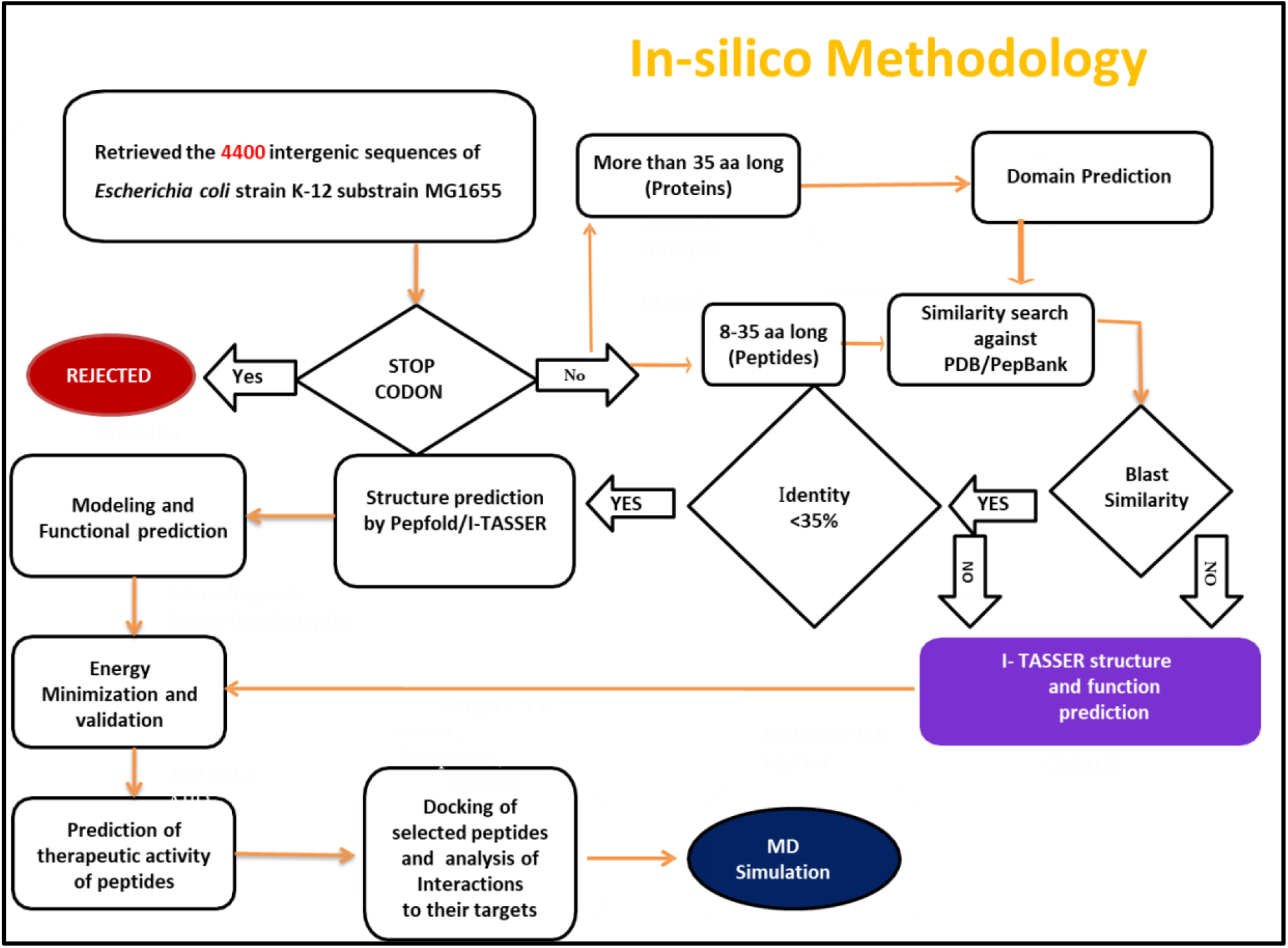
Bioinformatic workflow for database creation of ECOIs

**Fig 2:**
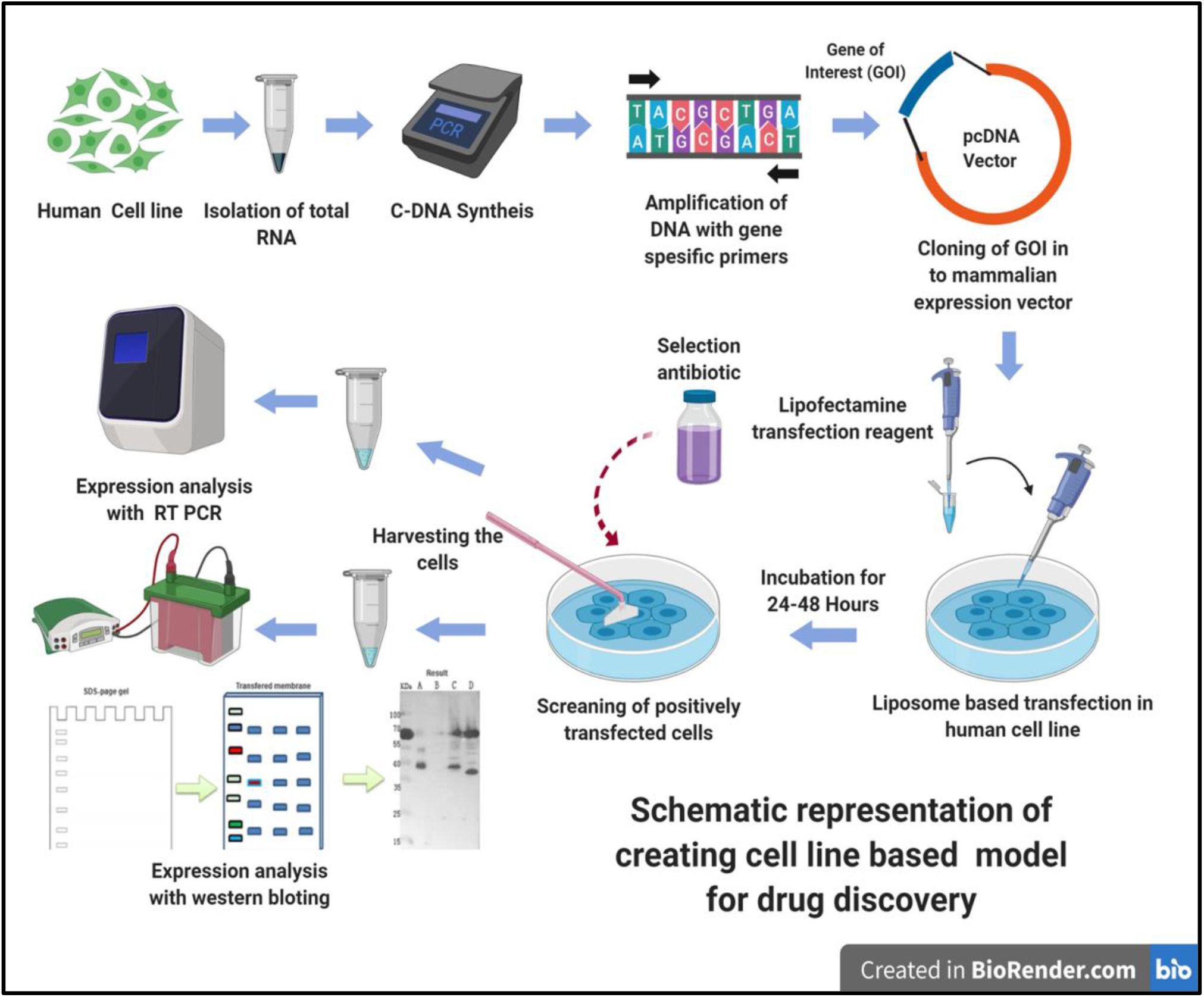
Methodology adapted for overexpression of target proteins in Human neuroblastoma Cell line (SH-SY5Y)

**Fig 3:**
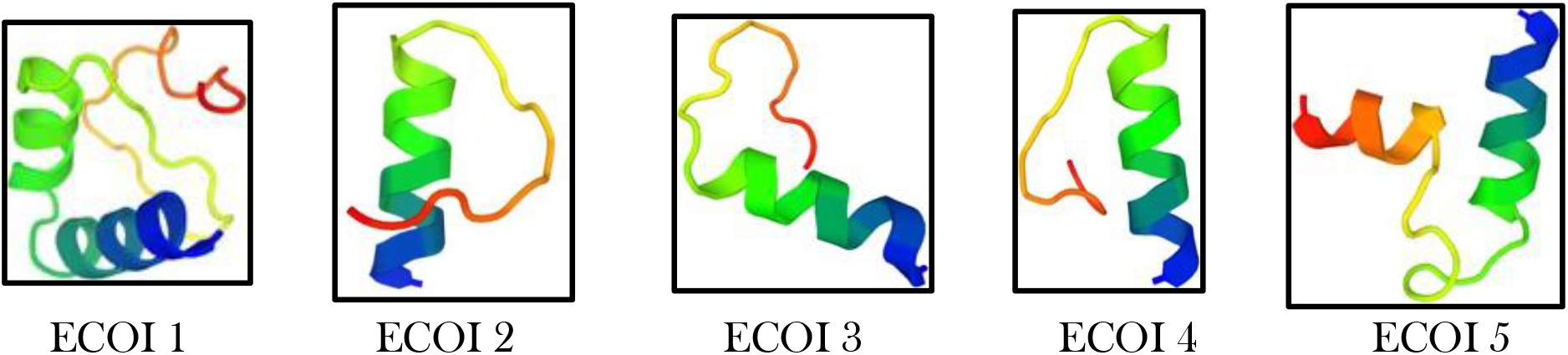
3 D structure of 5 lead ECOIs

**Fig 4:**
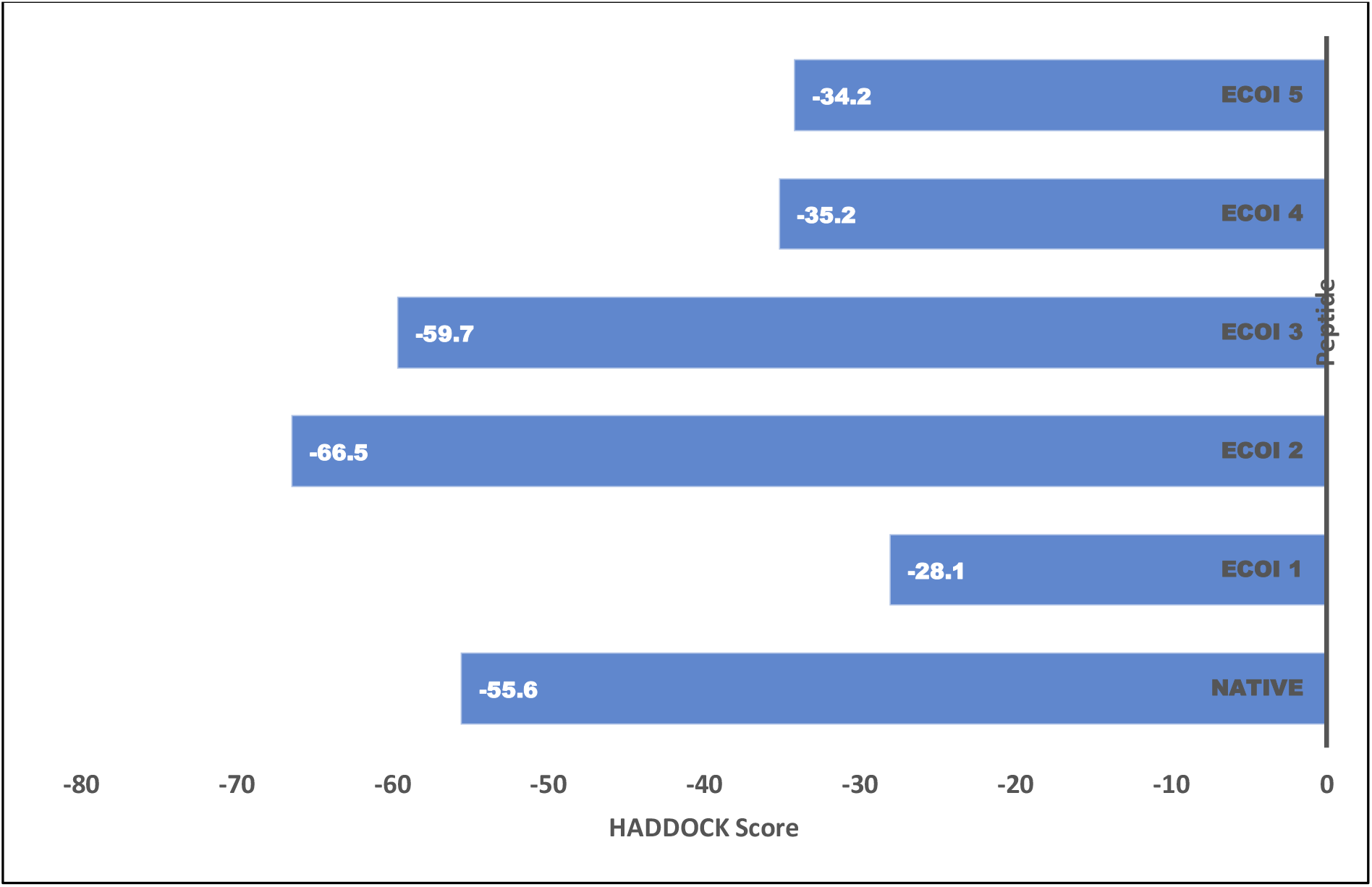
HADDOCK score of all the test and control peptides docked with BACE1 protein

### Docking study of the two most potent peptides (ECOI2 and ECOI3) against BACE1 target

Molecular docking of ECOI 2 and 3 was performed with human BACE1 protein as receptor (PDB Id 1FKN) using the HADDOCK2.4 server. Analysis of interacting amino acid residues of ECOI 2 and 3 showed strong binding to the active site of BACE1 (Fig. 5). The docking studies revealed that ECOI 2 and 3 make interactions with BACE1 active site residues at slightly different amino acids.

**Fig 5:**
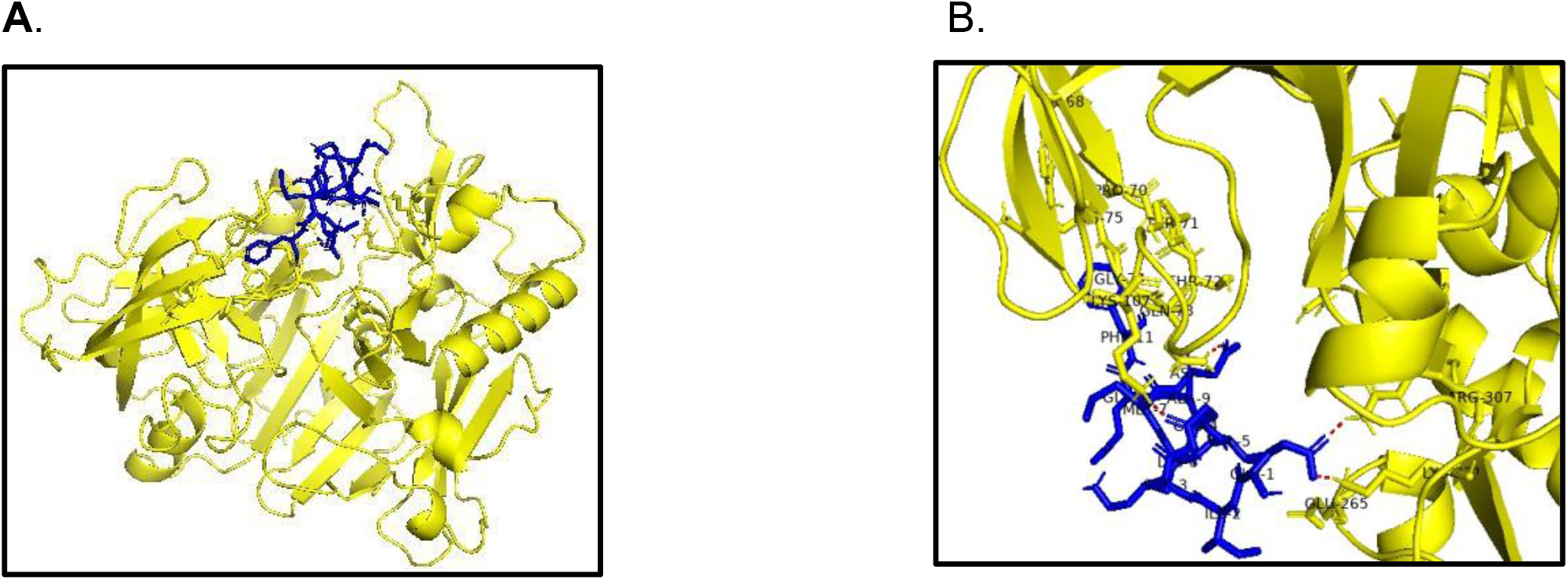
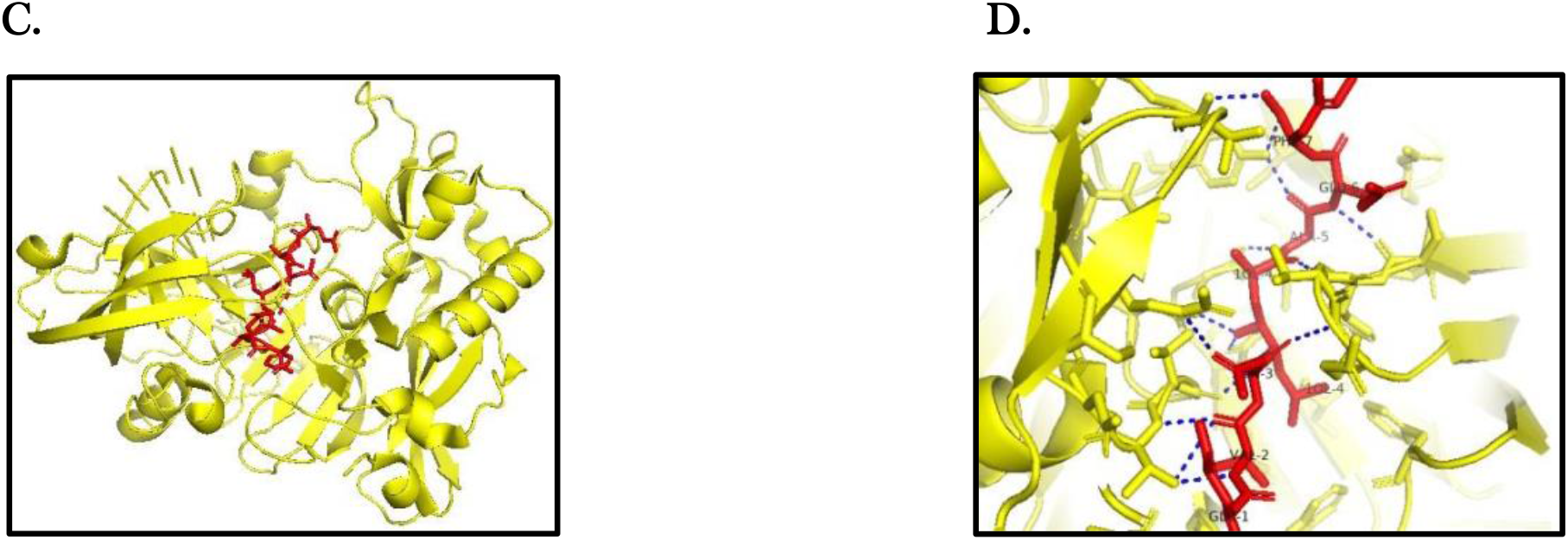
Docking of ECOI 2 and ECOI 3 with human BACE1 (PDB Id 1FKN) using HADDOCK2.4 server (A) Predicted binding mode of ECOI 3 (Blue) structures with BACE1 (Yellow). (B) Predicted binding mode of ECOI 2 (Red) structures with BACE1. (C) and (D) Molecular interactions between ECOI 2 and ECOI 3 with the pocket residues of the BACE1 active site. ECOI 2 and 3 established the important interactions with key pocket residues of the BACE1 active site (Asp32, Asp228) and the flap region (Tyr71, Thr 72, Gln 73, and Gly 74).

### In-vitro analysis of the effect of BACE1 inhibitory peptides (ECOIs)

As described in Table 2. ECOI 1,4 and 5 did not show a significant decrease in the activity of BACE1. But ECOI 2 and ECOI 3 led to a reduction in BACE enzyme activity up to 86.77 % ± 1.97 and 44.58 % ± 0.02 respectively. The activity inhibition of BACE1 was represented as the mean value of the percentage inhibition ± standard deviation.

**Table 1:**
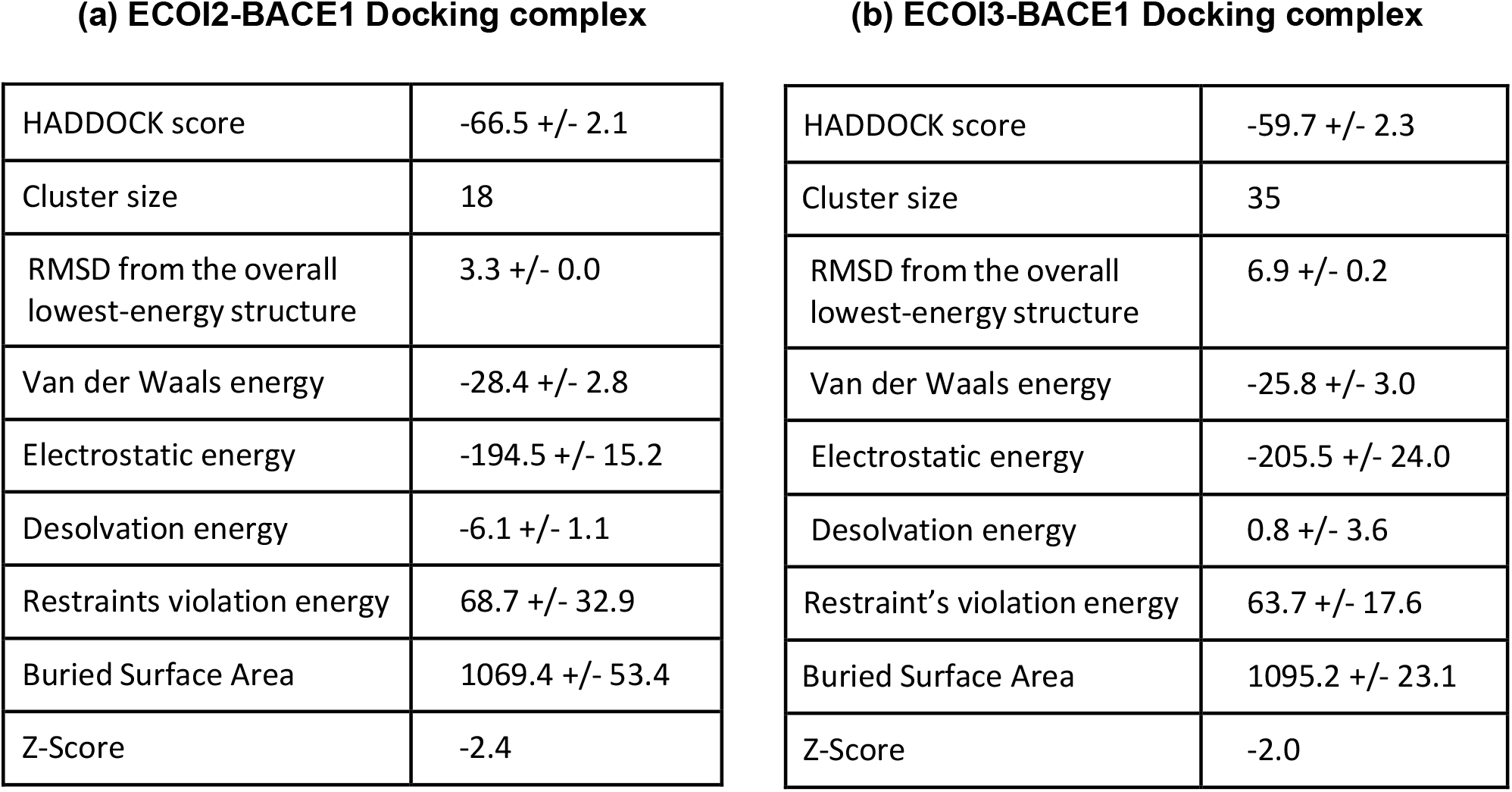
Docking complex (ECOI2-1FKN & ECOI3-1FKN) affinities scores predicted by HADDOCK

**(a) ECOI2-BACE1 Docking complex**
|  |  |
| --- | --- |
| HADDOCK score | -66.5 +/- 2.1 |
| Cluster size | 18 |
| RMSD from the overall lowest-energy structure | 3.3 +/- 0.0 |
| Van der Waals energy | -28.4 +/- 2.8 |
| Electrostatic energy | -194.5 +/- 15.2 |
| Desolvation energy | -6.1 +/- 1.1 |
| Restraints violation energy | 68.7 +/- 32.9 |
| Buried Surface Area | 1069.4 +/- 53.4 |
| Z-Score | -2.4 |

|  |  |
| --- | --- |
| HADDOCK score | -59.7 +/- 2.3 |
| Cluster size | 35 |
| RMSD from the overall lowest-energy structure | 6.9 +/- 0.2 |
| Van der Waals energy | -25.8 +/- 3.0 |
| Electrostatic energy | -205.5 +/- 24.0 |
| Desolvation energy | 0.8 +/- 3.6 |
| Restraint's violation energy | 63.7 +/- 17.6 |
| Buried Surface Area | 1095.2 +/- 23.1 |
| Z-Score | -2.0 |

**Table 2:**
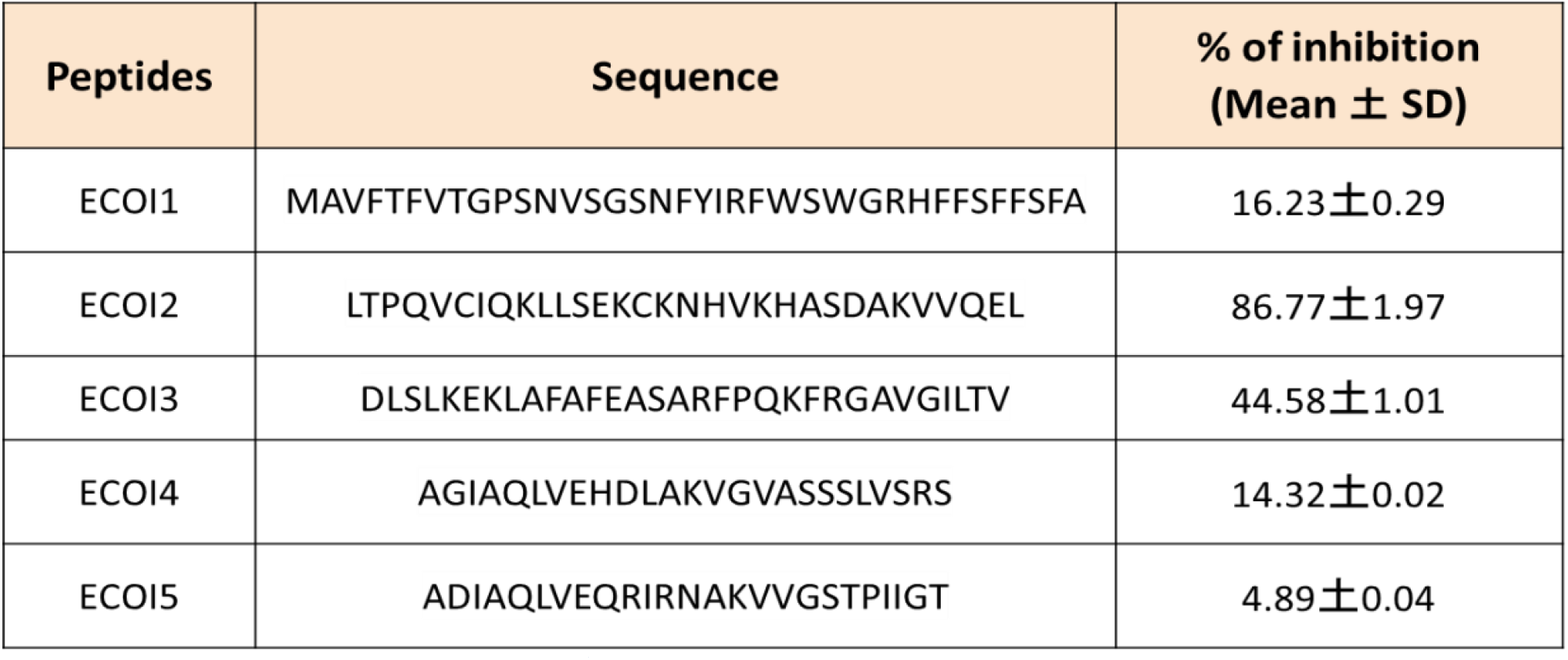
Amino acid sequences of the ECOIs (synthetic peptides). The results are presented as percentage activity inhibition of BACE1 and the average of three separate experiments.

### In-Vito BACE1 inhibition detection by FRET assay

The quantity of substrate cleavage was significantly reduced to 60.2% using a 100 nM ECOI2 concentration. On increasing the concentration of ECOI2 from 100 nM to 1000 nM, the amount of substrate cleaved was reduced from 65.97% to 86.77%. (Figure 6 A). Similarly, in the ECOI3 treated cells, the quantity of substrate cleavage was reduced to 40.2% using a 100 nM ECOI3 concentration. However, a non-significant change was observed when the concentration of ECOI3 was increased from 100 nM to 1000 nM (Figure 6B). The Beta secretase inhibitor IV (C_31_H_38_N_4_O_5_S) served as positive control in these experiments.

**Fig 6A and B:**
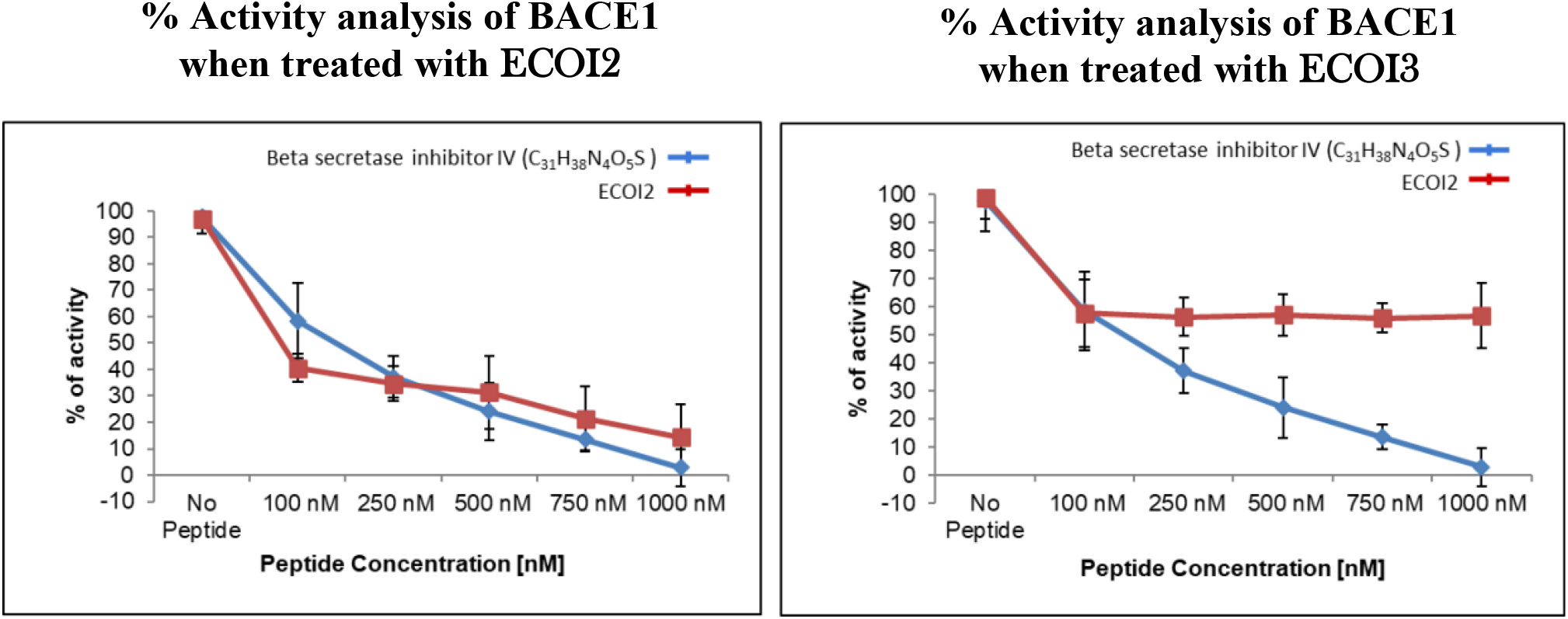
Percentage of activity analysis of BACE1 at different concentrations of ECOI2 and ECOI 3. Recombinant BACE1 (Supplied with kit) was pre-incubated with a concentration range of the peptides (100 nM-1000 nM) at 37°C for 4 hrs. in the dark followed by the addition of the fluorescently tagged substrate i.e., 7-Methoxycoumarin-4-acetyl-[Asn670, Leu671]-Amyloid /A4 Precursor Protein 770 Fragment 667–676-(2,4 dinitrophenyl) Lys-Arg-Arg amide trifluoroacetate salt. Fluorescence was measured using a 96-well plate in a multilabel plate reader (PerkinElmer, Waltham, MA, USA) using a wavelength of 320 nm excitation and 405 nm emission. Positive Control: Beta secretase inhibitor IV (C_31_H_38_N_4_O_5_S)

### ECOIs decrease Aβ 1-42 and Aβ 1-40 levels in Neuroblastoma SH-SY5Y cells

Results show that ECOI 2 below 250 nM did not affect Aβ1-40 and Aβ1-42 productions in SH-SY5Y-APP-BACE1 cells after 48 hours of treatments (Fig 7). The Aβ1-42 production was found to be reduced to 14.01±2.96% and 37.75±4.19% in presence of ECOI 2 and ECOI 3 respectively. Similarly, Aβ1-40 synthesis was found to be reduced to 9.09±6.86% and 33.86±6.19% by ECOI 2 and ECOI 3 of control respectively (ELISA). Results suggest that ECOI 2 profoundly affects Aβ metabolism in SH-SY5Y-APP-BACE1 cells.

**Fig 7A and B:**
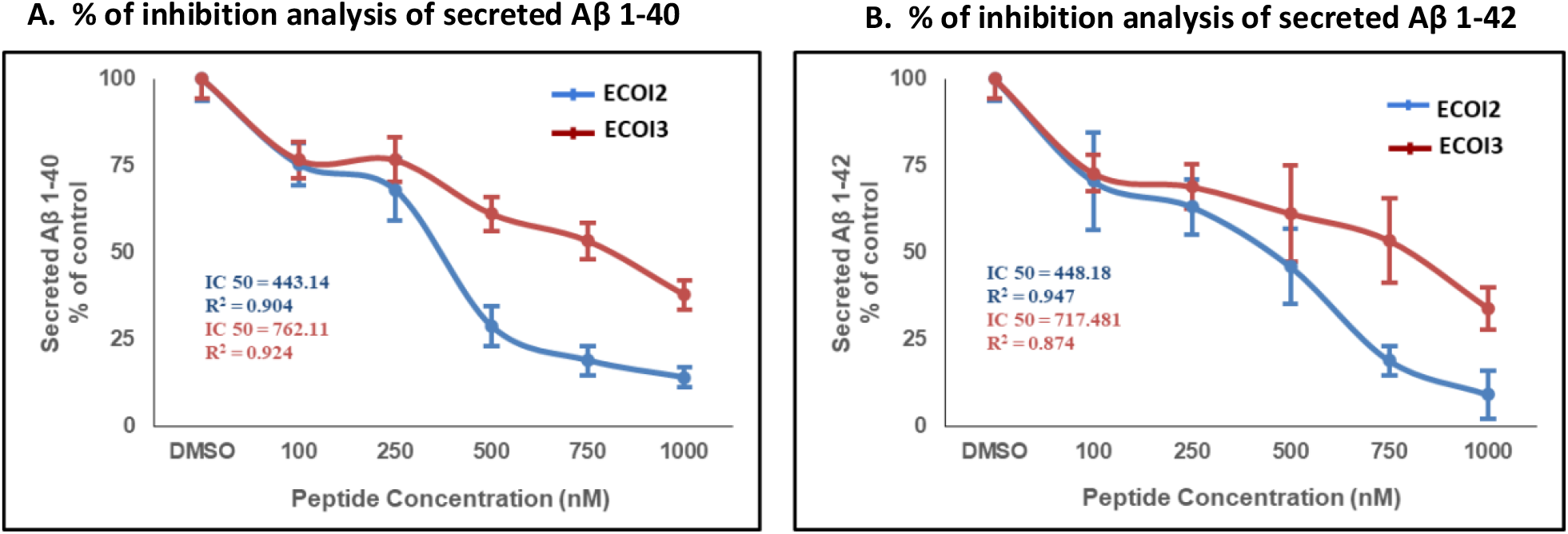
Percentage of inhibition analysis of BACE1 at different concentrations of ECOI2 and ECOI3. The BACE1 inhibitors decrease Aβ 1-40 and Aβ 1-42 levels in Neuroblastoma-SH-SY*5*Y cells constitutively expressing APP and BACE1. Cells were incubated in a serum-free medium with 100 to 1000 nM ECOI 2 or ECOI 3 for 24 h, at 37°C, in a humidified incubator with *5*% CO^2^. Cells with only DMSO (no peptides) were used as control. The levels of secreted Aβ 1-40 (A) and Aβ 1-42 (B) were determined by ELISA. The results are expressed as a percentage of control and represent tire mean± SEM of 3 independent experiments

**Fig 8:**
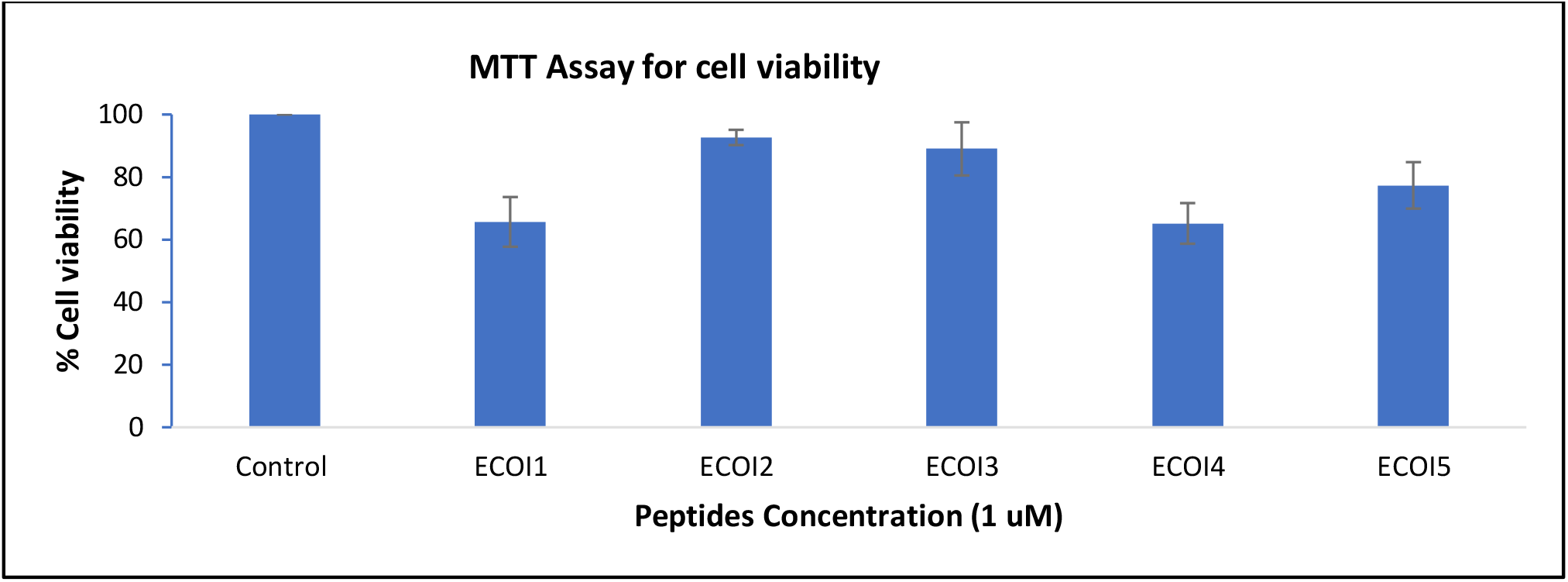
Cytotoxic assay of 5 lead peptides; 1.3×105 cells/cm2 are seeded. The new BACE1 inhibitory peptides ECOI 1-*5* at a concentration near the 1 uM. ECOI 1,4,*5* shows a significant decrease in cell viability. Whether ECOI 2 and 3 do not induce cytotoxicity at this concentration. SH-SY*5*Y-APP-BACE1 cells were incubated with 1 μM in an FBS-free culture medium for 24 h, at 37°C, in a humidified incubator with *5*% CO_2_ Untreated cells were used as control.

**Fig 9:**
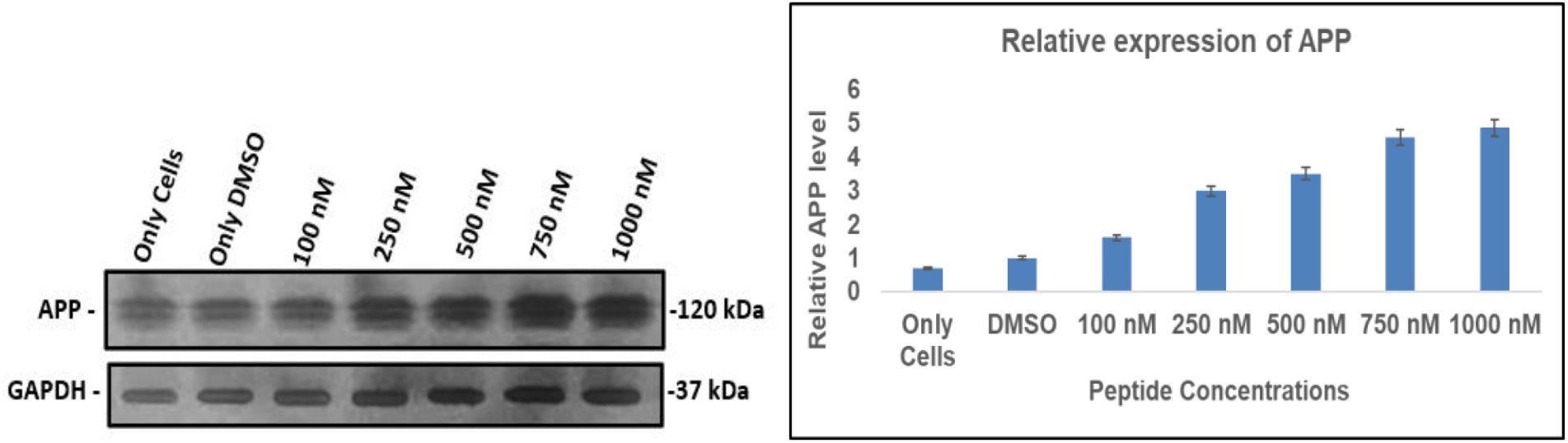
Evaluated AFP protein expression by western blot analysis after 24 hours of treatment with ECOI2 in the SH-SY*5*Y cell model. Only DMSO was regarded as the control group. Data showed that APP expression was significantly increased in a dose-dependent maimer in the ECOI2-treated cells than in non-treated SH-SY*5*Y. These results indicate that our cell model expresses high levels of APP due to inhibition of BACE1 activity.

### Cytotoxicity effect of peptides lead (MTT Assay)

The MTT assay of ECOIs showed non-significant changes in the proliferation rate of SH-SY5Y stable cells in the presence of ECOI 2 and 3. The ECOI 1,3, and 5 showed a 30-40% reduction in the cell proliferation rate. Although 100% cell viability was reported in the control reaction (i.e., no peptide), ECOI 2 and 3 were the lead peptides for further experiments due to their non-cytotoxic effect.

### The ECOI 2 selectively inhibit the APP cleavage

The relative expression of APP protein has been analyzed with or without the presence of ECOI2 at different concentrations (100 nM-1 μM). It has been observed that there were no significant changes occurred when cells treated with DMSO and 100 nM of peptide concentration as compared with the cells without treatment.

As the concentration of ECOI2 was increased from 100 nM to 1 μM the accumulation of APP protein was significantly increased, suggesting that APP accumulation occurred due to the inhibitory effect of the BACE1 enzyme.

### ECOI2 inhibits the BACE1 expression

The ECOI2 peptide was considered as a prominent lead due to its ability to decrease the Aβ levels (Figure 6A). Western blotting results suggested that a decrease in the expression of BACE1 protein on treating the cells with ECOI2 at a concentration of 500 nM and higher. The expression of BACE1 was relatively decreased with increasing concentrations of ECOI2, validating the hypothesis that BACE1 expression was decreased along with the activity of BACE1. The western blot results suggested that the ECOI2 is responsible for activity inhibition and also the regulation of the expression of the BACE1 in an in-vivo system.

### Effects of ECOI 2 on the mRNA Levels of Aβ generation related genes

To examine the effect of ECOI2 on Aβ generation-related genes, the mRNA levels of most genes examined, like PS1, APOE, LRP1 and LDLR, remained unchanged by the treatment of ECOI 2 compared with SH-SY5Y-APP-BACE1 only cells at 12-hour (Fig. 11). However, with a 12-hour ECOI 2 treatment, gene expression of PS2 was significantly up-regulated, whereas expression IDE of was down-regulated compared with SH-SY5Y-APP-BACE1 only cells; Similarly, the expression of the APP gene was increased when treated with ECOI 2 whereas the expression of BACE1 is downregulated as supported by western blot analysis (Fig. 11).

**Fig 10:**
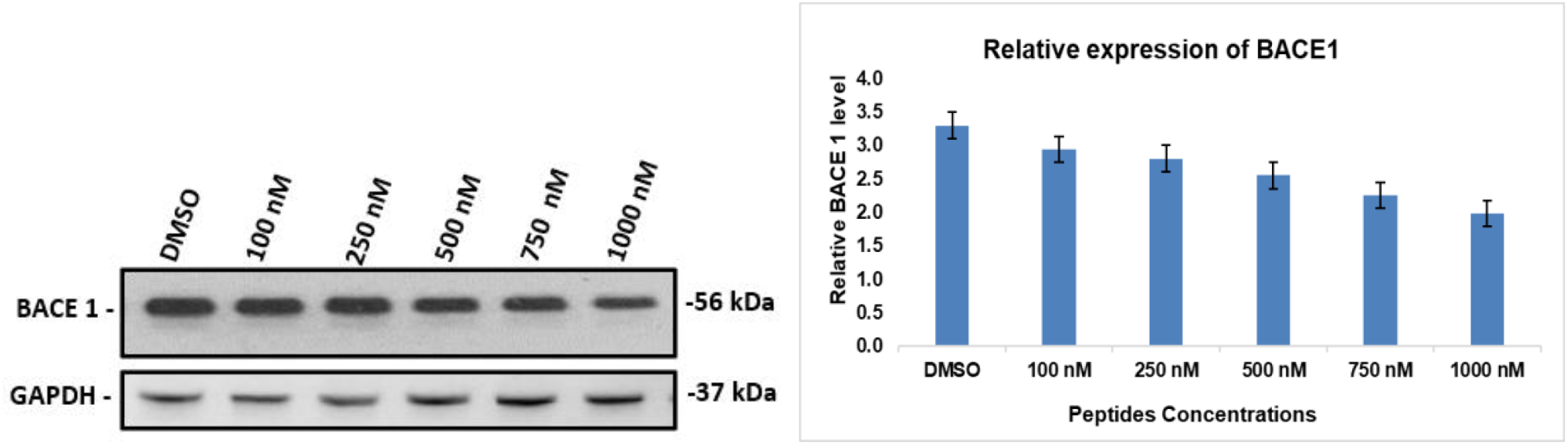
BACE1 protein expression evaluation by western blot analysis after 24 hours of treatment with ECOI2 in the SH-SY*5*Y cell model. Only DMSO was regarded as the negative control. Data showed that BACE1 expression was significantly reduced compared to non-treated SH-SY*5*Y cells. Results indicate that the cell model successfully expressed high levels of BACE1 showing reduction in die expression of BACE1 protein in the presence of ECOI2.

**Fig 11:**
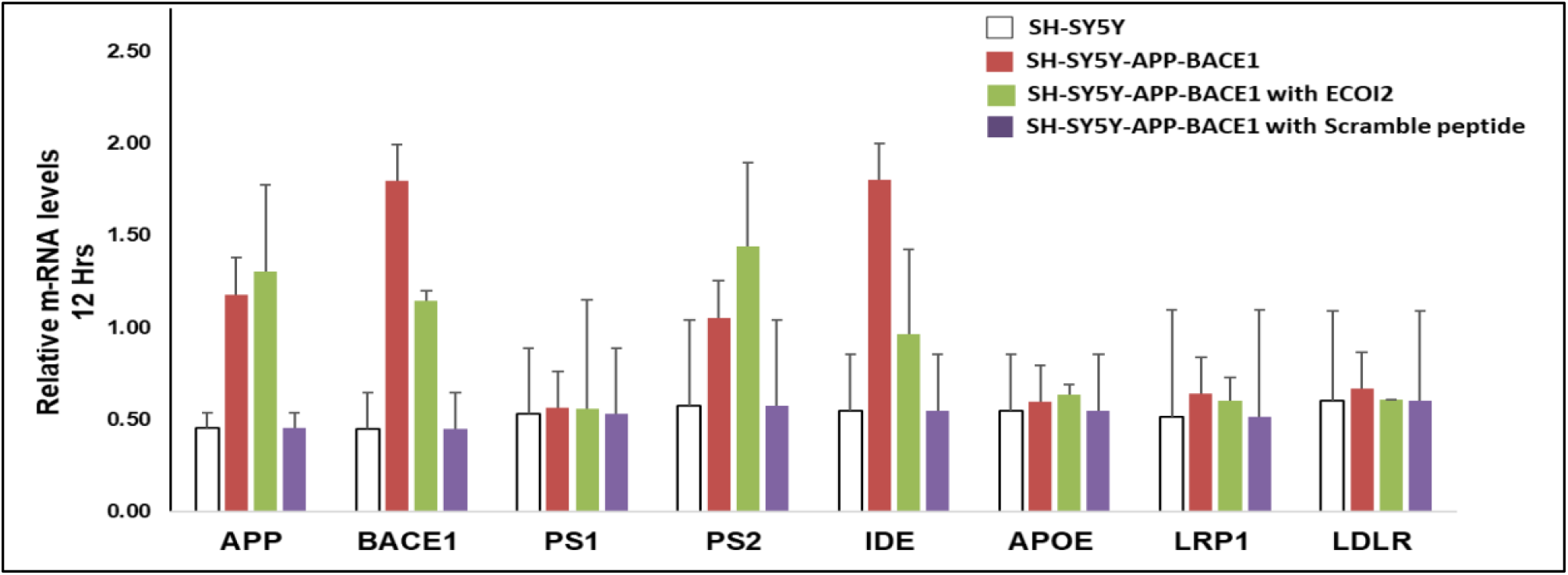
Effects of ECOI 2 on the mRNA levels of Aβ generation-related genes. The relative mRNA expression levels were determined by real-time PCR. APP, BACE1, PS1, PS2, IDE, APOE, LRP1, and LDLR after treatment with 1000 nM of ECOI 2 for 12 hours. Non-treated SH-SY5Y cells were regarded as a control and SH-SY5Y-APP-BACE1 treated scramble peptide of ECOI 2 was considered as a negative control. Data are expressed as the mean ± SD of duplicate reaction.

The expression of BACE1 and APP genes in SH-SY*5*Y-APP-BACE1 cells was found to be upregulated when compared with only SH-SY*5*Y cells (non-transfected). The data supported the overexpression of APP and BACE1 on transfection and confirmed the stable expression of APP and BACE. The scrambled peptide of ECOI 2 served as a negative control with the same amino acid composition but different order of amino acid sequence.

## Discussion

This study was designed to find new therapeutic peptides from the non-expressed intergenic sequences of Escherichia coli, which make we what call as the dark matter of the genome.

The term ‘dark genome’ includes the non-expressing, non-translating, and extinct DNA sequences that can be artificially encoded into functional molecules. The non-expressing sequences include antisense, reverse coding, repetitive sequences, intergenic sequences. The non-translating sequences include tRNA, ncRNA, ribosomal RNA, and introns. The extinct component of the dark genome includes pseudogenes that were functional at one time but are no longer active.

The first objective of this study was to narrow down a huge search space in the dark genome of E.coli and create a list of potential therapeutic peptides that show anti-Alzheimer property. Towards this goal, 4400 E. Coli K12 MG1655 intergenic sequences were computationally studied for their uniqueness and efficacy. The search for dark peptides was restricted to 10-35 residues based on the ability to synthesize these peptides. Their physicochemical, structure and function properties were computationally studied and the topmost peptides were marked for experimental validation.

Results suggested that ECOI2 and ECOI3 were promising peptide candidates for the development of the first-in-the-class anti-Alzheimer drugs. These peptides strongly inhibited the activity of the BACE1 enzyme and reduced Aβ accumulation in vitro. Furthermore, peptides showed no cytotoxic effect. The scrambled peptide of ECOI2 did not change the mRNA level of expression, suggesting that a certain order of amino acid sequences was responsible for the observed activity.

Previous reports on molecular determinants of Alzheimer’s have demonstrated the effect of synthetic peptides by blocking the active site of the BACE1 enzyme - a key drug target in Alzheimer’s disease (Resende et al., 2020). The royal jelly peptides isolated from natural sources have also been utilized as potential inhibitors of BACE1 (Xueqing Zhang et al. 2019). These studies encouraged us to look for novel genomic spaces in search of anti-BACE1 candidates.

Following a series of bioinformatics steps, the two most promising peptides (ECOI2 and ECOI3) were shortlisted for experimental evaluation. The in-vitro studies with recombinant BACE1 strongly suggest that ECOI 2 and ECOI 3 inhibit the BACE1 enzyme activity and were found to reduce the cleavage of APP. Furthermore, the cell-penetrating property and lack of cytotoxic effects enhanced their pharmacological potential.

The impact of ECOI2 and ECOI3 peptides was determined using the in-vitro and in cell-based systems leading to a significant reduction in Aβ40 and Aβ42 accumulation without cytotoxic effects. Using the BACE1 and APP sequences cloned in the mammalian expression vector, a cell-based Alzheimer’s disease model was created to investigate the therapeutic potential of proposed anti-Alzheimer molecules.

Experimental evidence clearly indicate that ECOI2 and ECOI3 peptides decreased Aβ levels by inhibiting the activity of the BACE1 enzyme in the cell-based Alzheimer’s disease model. The BACE1 inhibitor (ECOI2 and 3) showed a significant quantitative decrease in the SH-SY5Y cells after treatment with these peptides in the concentration range of 100nM -1000nM.

Though the harvesting of novel therapeutic peptides against Alzheimer’s are encouraging, this is an initial proof of concept using a limited number of in-vitro possibilities. It would be interesting to confirm the results in suitable animal models. Furthermore, the study evaluated the effect of ECOI2 and ECOI3 on Aβ accumulation. It is quite possible that the peptides may have result in other biochemical and physiological correlations beyond the scope of this study.

Here, we investigated the effect of ECOI2 on the expression of genes related to Aβ generation. Upon ECOI2 treatment of the cells, a significant up-regulation of PS2 gene expression and a down-regulation of IDE gene expression was unexpectedly observed.

In future, this study needs to expand to identify molecular mechanisms of ECOI2 and ECOI3 and extend the study in suitable animal models of Alzheimer’s disease.

Beyond the therapeutic potential of ECOI peptides, this work leads to an important realization that dark genome is a huge untapped goldmine that is waiting for an exploration by pharma as a novel drug discovery platform.

## Acknowledgements

This work is supported by DBT JRF Fellowship (DBT/2016/JNU/741). We express our sincere gratitude to Dr Neha Channa for providing vectors used in this study.

## Author contribution

NV performed all the computations and experiments and wrote the first draft of the paper. SM supervised the experimental work. PKD conceived the idea, designed the study, and finalized the manuscript

